# Dynasore, the dynamin inhibitor, modulates longitudinal bone growth in a hormetic manner

**DOI:** 10.64898/2026.03.09.709778

**Authors:** Jose G. Marchan-Alvarez, Sanya Koikkara, Ruihan Zhou, Oscar P. B. Wiklander, Phillip T. Newton

## Abstract

**Background:** Longitudinal bone growth occurs via the process of endochondral ossification, involving a complex interplay of chondrocyte proliferation, differentiation, and matrix remodelling. As with all mammalian cells, chondrocytes require dynamin for mitochondrial fission, to shuttle vesicles from the Golgi apparatus, and for both clathrin- and caveolin-mediated endocytosis. Here, we aimed to test the functions of dynamin on bone growth. To do so, we applied dynasore - a small molecule that is a reversible dynamin inhibitor - to mouse metatarsal bones cultured *ex vivo*. We assessed gross changes using bone length measurements and histomorphometry, and combined this with EdU detection, immunostaining, super-resolution microscopy and transmission electron microscopy.

**Results:** Dynasore induced a dose-dependent hormetic effect on bone elongation: while high concentrations (220 µM) impaired growth and abolished chondrocyte proliferation, low-dose treatment (40 µM) significantly increased longitudinal bone growth. Histological analysis demonstrated that low dose dynasore augmented epiphyseal cartilage expansion and matrix accumulation, particularly within the resting and proliferative zones, while reducing chondrocyte proliferation. Immunostaining indicated that 40 µM dynasore preserved collagen type X synthesis, activated mTORC1 signalling, and blocked autophagy, based on SQSTM1 accumulation. Low dose dynasore treatment expanded the thickness of the filamentous actin layer at the plasma membrane and deepened collagen fiber-containing endocytic pits, indicating that impaired cartilage remodelling was associated with growth-associated matrix accumulation.

**Conclusions:** This study reveals that dynasore exerts hormetic effects on growth plate chondrocytes, wherein low doses stimulate bone elongation, and high doses impair chondrocyte function.

## 1. Introduction

During childhood and adolescence, humans grow taller due to the elongation of bones such as those in our legs and spine. These bones grow through a process called endochondral ossification, whereby chondroprogenitor cells undergo sequential differentiation into proliferative and hypertrophic chondrocytes, which remodel and mineralise their hyaline cartilage extracellular matrix (ECM). Mineralised cartilage is then used as a scaffold on which new bone tissue is deposited by osteoblasts, ultimately contributing to bone elongation and height gain in children and adolescents [1].

The protein dynamin is crucial for certain forms of membrane scission that are conserved among higher eukaryotes [2]. Dynamin is a GTPase that is recruited into oligomeric helices that form around a lipid bilayer membrane tube. The dynamin oligomer then undergoes conformational changes that constricts the membrane tube, narrowing it to a point where the inner luminal radius falls below 2 nm, prior to fission of the membrane in a GTP-dependent manner [3]. Thus, dynamin is required for membrane trafficking processes, such as clathrin- and caveolin-mediated endocytosis [4], shuttling of vesicles from the Golgi apparatus as well as mitochondrial fission [5]. While its functions mean that dynamin is involved in a large number of cellular processes, our understanding of its roles in chondrocytes during bone growth is incomplete.

Dynasore is a cell-permeable small molecule that acts as a non-competitive, irreversible inhibitor of dynamin. By binding to the GTP-binding domain, dynasore specifically targets the GTPase activity of dynamin, thereby preventing the hydrolysis of GTP required for membrane fission [4]. The actions of dynasore in disrupting the scission of vesicles at the inner leaflet of the plasma membrane [6] (Fig. 1A), has been observed in different cell types such as immune cells [7], muscle cells [8], osteosarcoma cells [8], and articular chondrocytes [9]. Yet, beyond its classical role as a dynamin inhibitor, dynasore may alter other processes in a dynamin-independent manner, including cholesterol homeostasis, lipid raft integrity, lysosomal inhibition[10] and actin remodelling [5]. Whereas the properties of dynasore have been explored in the articular cartilage [9], the effects on endochondral ossification remain to be determined.

**Fig. 1.**
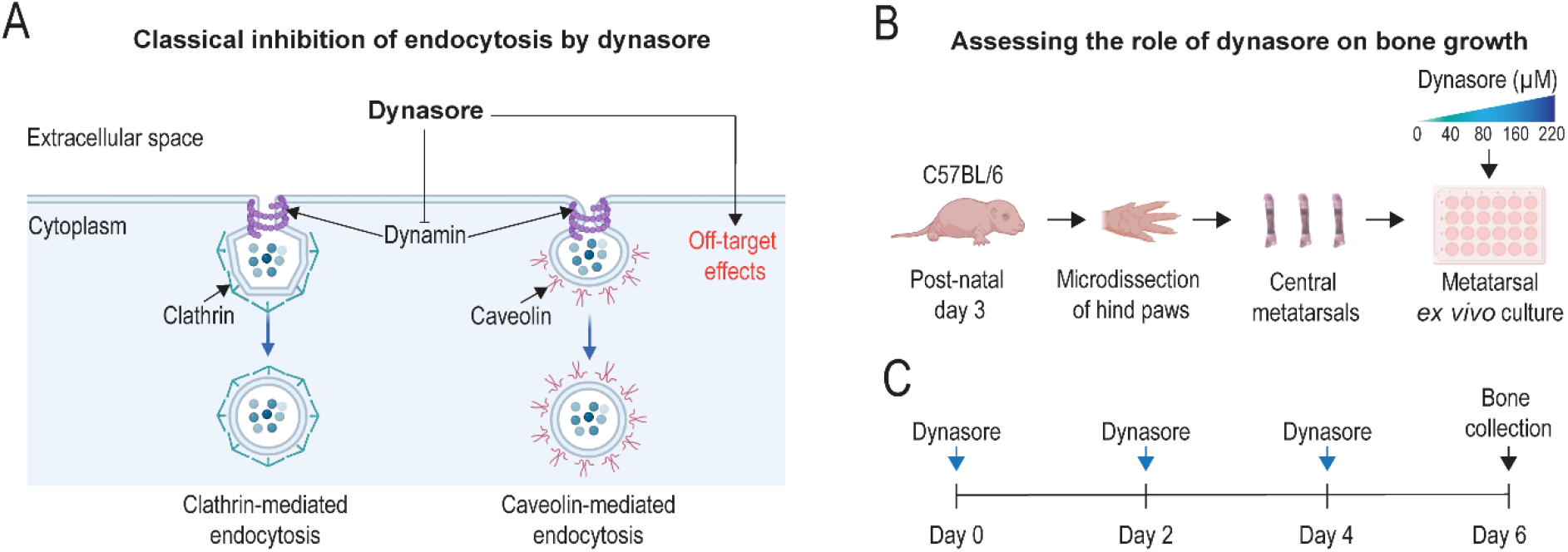
Experimental workflow. **A** Schematic overview of dynasore functions; whilst dynasore has been widely used as a dynamin inhibitor to block various endocytic pathways, including clathrin-mediated and caveolin-mediated endocytosis, less well described dynamin-independent effects are also reported. **B, C** Central metatarsals were isolated from the hind paws of C57BL/6 mice and cultured *ex vivo*. Dynasore was then applied in a dose-response manner at days 0, 2 and 4.

Hence, we aimed to test the effects of dynamin inhibition mediated by dynasore on longitudinal bone growth.

## 2. Materials and Methods

### 2.1. Ethical Approval

All experiments in this study were approved and performed in accordance with the guidelines from the Swedish National Board for Laboratory Animals and the European Union Directive (2010/63/EU), using an ethical permit (DNR 16673/2020) approved by Stockholm’s Animal Experimentation Ethics Committee (Stockholms djurförsöksetiska nämnd). The animals were purchased from Charles River (strain code: 632).

#### 2.2. *Ex vivo* culture of murine metatarsals

This system allows the study of chondrocytes at broad differentiation statuses within their intact three-dimensional tissue architecture, providing a more physiologically relevant model compared to monolayer or primary chondrocyte cultures [11–13]. Under a stereo-microscope (Nikon SMZ645) and aseptically, the three central metatarsals were microdissected from both hind paws of three-day-old C57BL/6 mice. Microdissection was performed in pre-cooled (4 °C) DMEM/F-12 (Dulbecco’s Modified Eagle Medium/Nutrient Mixture F-12) with GlutaMAX™ (Gibco™, catalog number: 31331093) and supplemented with 50 mg/ml gentamycin (Gibco™, catalog number: 15750045). Metatarsals were cultured for up to six days in DMEM/F-12 medium supplemented with 10 mM β-glycerophosphate (BGP) (Sigma-Aldrich, catalog number: G9422), 50 µg/ml L-ascorbate-2-phosphate (ascorbic acid) (Sigma-Aldrich, catalog number: G5960),0.2% bovine serum albumin (Sigma-Aldrich, catalog number: 05470) and 20 mg/ml gentamycin, as previously reported [13]. Bone cultures were maintained in an incubator (37 °C, 5% CO_2_) and the medium was changed every two days. Three central metatarsals micro-dissected from one paw were used as controls (vehicle-treated) whereas the three central metatarsals from the contralateral paw were used as the experimental group (treatment). Therefore, each observation is the average of all three bones from one paw (i.e. one observation = one animal), unless otherwise stated in the figure legend.

#### 2.3. Metatarsal treatment with dynasore and EdU

Dynasore (Sigma-Aldrich, catalog number: 324410) was prepared as a 20 mM stock solution in DMSO (Sigma-Aldrich, catalog number: D8418), aliquoted into 20 µl portions, and stored at -80 °C in the dark until further use. *Ex vivo* metatarsal cultures, as described in section 2.1, were treated with dynasore (at concentrations of 40 µM, 80 µM, 160 µM, and 220 µM) on days 0, 2, and 4. Bafilomycin (8 nM; Thomas Scientific, catalog number: C974N94), a well-established compound known to induce overgrowth in tubular bones [13], was used as a positive control, and also used in combination with 40 µM dynasore. EdU (ThermoFisher, catalog number: A20012) was prepared in distilled water as a 5 mM stock solution and kept at - 20 °C. Subsequently, EdU at 5 µM was added to the metatarsal cultures 4 hours before collecting the bones at day 6.

Longitudinal bone growth was documented every two days (days 0, 2, 4, and 6) using an Infinity 1 camera integrated with a Nikon SMZ-U stereo-microscope. Bone length measurements were subsequently performed using ImageJ analysis software (National Institutes of Health).

#### 2.4. Safranin O/Fast Green staining of metatarsals

Metatarsals were fixed in 4% paraformaldehyde (PFA) for 48 hours, then transferred to 10% ethylenediaminetetraacetic acid (EDTA, pH 8.05; Scharlau, catalog number: AC09401000) for 48 hours followed by 70% ethanol (EtOH). These incubations were carried out under gentle shaking on a rotating platform (IKA, KS-130 basic) at 4 °C. Next, bones underwent dehydration overnight in a LOGOS Microwave Hybrid Tissue Processor (Milestone). The dehydrated samples were embedded in paraffin and sectioned at a thickness of 5 µm using a microtome (Marshall Scientific, Microm HM 360). The 5 µm sections were floated on heated water (60 °C) to spread them out and then mounted on plus gold slides (ThermoFisher, catalog number: 11976299). Thereafter, bone sections were incubated at 60 °C for 1 hour in a Hybaid oven to remove paraffin, followed by sequential rehydration through xylene (twice), 99% EtOH, 95% EtOH, and 70% EtOH, with each step lasting 5 minutes. The samples were then stained with Weigert’s Hematoxylin, prepared with hematoxylin (Sigma-Aldrich, catalog number: H3136) and ferric chloride (Sigma-Aldrich, catalog number: 157740), for 5 minutes, rinsed in distilled water for 3 minutes, briefly dipped (2 seconds) in acid alcohol, and washed again with distilled water. Next, the slides were stained with 0.02% Fast Green (Sigma-Aldrich, catalog number: F7258) for 3 minutes, treated with 1% acetic acid for 3 minutes, and, without rinsing, stained with Safranin-O solution (Sigma-Aldrich, catalog number: S8884) for 30 minutes. Following staining, the samples were rinsed in 95% EtOH and dehydrated through sequential steps of 95% EtOH, 99% EtOH (twice), and xylene (twice), each step lasting 5 minutes. Finally, the sections were mounted using Pertex mounting medium (Histolab, catalog number: 00811) and imaged using a Zeiss AX10 microscope equipped with an Axiocam MRm camera.

Histomorphometry analyses of the growth plate zones and hypertrophic cell height were performed using ImageJ. The relative amount of matrix was also quantified using ImageJ software. Briefly, images were converted to 8-bit format, the epiphyseal cartilage boundary (including either “RZ+PZ” or “HZ”) was delineated, and a Gaussian blur filter (radius = 2) was applied before intensity thresholding. The matrix content was then calculated as the percentage of the total area.

#### 2.5. Multiplex immunofluorescence for phospho-RPS6 and SQSTM1

Bone sections were incubated at 60 °C to remove paraffin and then rehydrated following the protocol outlined in section 2.4. Next, the specimens were placed in DAKO target retrieval buffer (Agilent technologies, catalog number: S1699) and heated in a sealed pressure cooker (CertoClav A-4050) at 0.2 bar. After cooling the pressure cooker for 30 minutes, the samples were removed and rinsed three times with distilled water and washed once with phosphate buffered saline (PBS, 1X) containing 0.05% Tween 20 (PBST; Sigma-Aldrich, catalog number: p9416), hereafter known as PBST. Blocking was performed using 3% normal horse serum (Jackson ImmunoResearch, catalog number: 8000121) diluted in PBST with 0.05% Triton 100X (Sigma-Aldrich, catalog number: T8787) for 1 hour at room temperature. Primary antibodies targeting phospho-RPS6 (1:100; Cell Signaling, catalog number: 4858) and SQSTM1 (1:500; Progen, catalog number: GP62-C) were diluted in the blocking buffer and incubated with the samples overnight at 4 °C. The following day, the slides were equilibrated to room temperature for 15 minutes and washed five times with PBST, each wash lasting 5 minutes. Secondary antibodies (1:400) were prepared in blocking buffer and applied to the samples for 1 hour at room temperature. For phospho-RPS6, the secondary antibody was purchased from Invitrogen (catalog number: A31573), and for SQSTM1, it was acquired from Jackson ImmunoResearch Laboratories (catalog number: 706-545-148). After secondary antibody incubation, slides were rinsed once with PBST for 5 minutes. Subsequently, the sections were stained with DAPI (1.5 µg/ml; Sigma-Aldrich, catalog number: D9542) diluted in PBS (1X) for 20 minutes, followed by three washes with PBS (1X). The samples were mounted using Fluoroshield mounting medium (Sigma-Aldrich, catalog number: F6182) and imaged using a Leica Stellaris 5 X advanced line scanning confocal microscope. Quantification of positive cells for phospho-RPS6 and SQSTM1, relative to DAPI positive cells, were performed using QuPath software (version 0.5.1) [14].

#### 2.6. Immunostaining for collagen type X multiplexed with EdU staining

Bone were fixed in 4% PFA for six hours, followed overnight incubation in 30% sucrose solution (diluted in 1X PBS) (AG Scientific, catalog number: S-2885) on a rotating platform at 4 °C. The following day, the bones were embedded in optimal cutting temperature (OCT) medium (Sakura Finetek, catalog number: 4583) and frozen at -20 °C. Cryosectioning was performed at a thickness of 20 µm using an NX70 Cryostat (Epredia) and sections were placed on Plus Gold slides. The samples were air-dried and stored at -20 °C until further use. Prior to staining, the sections were thawed at room temperature and air-dried. The samples were then incubated in 1X PBS for 15 minutes in a humidified chamber. Subsequently, blocking was performed using 3% normal horse serum diluted in PBST containing 0.25% Triton X-100. Collagen type X primary antibody (1:750; Abcam, catalog number: 58632) was diluted in the blocking buffer and incubated with the samples overnight at 4 °C. Subsequently, the slides were washed three times with PBST for five minutes each, incubated with a secondary antibody (1:600; Invitrogen, catalog number: A31572) at room temperature and washed three times with PBST, each wash lasting 5 minutes. Simultaneously, the EdU reaction mix was prepared in a chemical hood. This mix contained 1 M TRIS (pH 7.5), 100 mM CuSO_4_, Alexa-Azide 647 (diluted 1:10) (Thermo Fisher Scientific, catalog number: A10277), distilled water, and 0.5 M ascorbic acid (diluted in distilled water). Samples were incubated with the reaction mix for 30 minutes, followed by a five-minute rinse with PBST. Nuclei were counterstained with DAPI for 15 minutes, and the samples were mounted using Fluoroshield, as described in section 2.5. Imaging was performed using a Leica Stellaris 5 X confocal microscope and quantifications of EdU positive cells (relative to DAPI) and length of collagen type X were processed with QuPath software.

#### 2.7. F-actin staining and super-resolution microscopy

Metatarsals were cultured under vehicle, bafilomycin or dynasore (40 µM) and processed for fixation, OCT embedding and cryo-sectioning as in 2.3 and 2.6 respectively. Then, tissue sections were incubated with 1X PBS for 15 minutes in a humidified chamber. Subsequently, bones were incubated simultaneously with Alexa Fluor 647–conjugated phalloidin (Thermo Fisher Scientific, Cat. No. A22287) to stain f-actin, and DAPI to label nuclei, for 45 minutes at room temperature. After staining, samples were rinsed three times with 1× PBS and mounted using Fluoroshield. Fluorescence images were acquired using a Zeiss LSM 980 confocal microscope equipped with an Airyscan 2 detector. Sequential imaging was performed using appropriate laser lines and filter settings for Alexa Fluor 647 and DAPI. Airyscan super-resolution mode was used where indicated.

Images were processed using ImageJ to measure the thickness of the f-actin staining around the inner leaflet of the plasma membrane, based on a 2D line [15]. Briefly, images were thresholded and converted to mask. Then a line was drawn using the longest edge containing f-actin. The width of f-actin staining was measured at each end of the line. Thus, two measurements were collected per cell, which were subsequently averaged.

#### 2.8. Transmission electron microscopy (TEM)

Metatarsals were culture under vehicle, bafilomycin or dynasore (40 µM) as in 2.3. Bone specimens were promptly fixed in a solution containing 2% glutaraldehyde and 1% paraformaldehyde, prepared in 0.1 M sodium cacodylate buffer supplemented with 0.1 M sucrose and 3 mM CaCl_2_ (pH 7.4), for 30 minutes at room temperature. Following fixation, samples were stored at 4 °C until further processing. The metatarsals were then rinsed in 0.1 M sodium phosphate buffer (pH 7.4) and post-fixed in 2% osmium tetroxide in the same buffer at 4 °C for 2 hours. Dehydration was carried out through a graded ethanol and acetone series, after which the specimens were embedded in LX-112 epoxy resin (Ladd Research). Ultrathin sections (∼80–100 nm) were prepared using a Leica EM UC7 ultramicrotome, stained sequentially with uranyl acetate and lead citrate, and imaged using a Hitachi HT7700 transmission electron microscope operating at 80 kV. Digital micrographs were captured with a 2k × 2k Veleta CCD camera (Olympus Soft Imaging Solutions).

For morphometric analysis, only cells with a fully visible perimeter in the electron micrographs were included as previously reported [16]. Briefly, all coated structures (shallow, U-shaped and Ω-shaped) located within 500 nm of the plasma membrane, were identified and quantified in the resting, proliferative, pre-hypertrophic and hypertrophic zones.

#### 2.9. Statistics and reproducibility

To increase the robustness and reproducibility of the data, every observation within each treatment group derives from a mouse obtained from a different litter, unless otherwise stated in the figure legend. Length measurements (Fig. 2, Fig S1, Fig. S2) and histomorphometry (Fig. 3) were performed in a single-blinded manner.

**Fig. 2.**
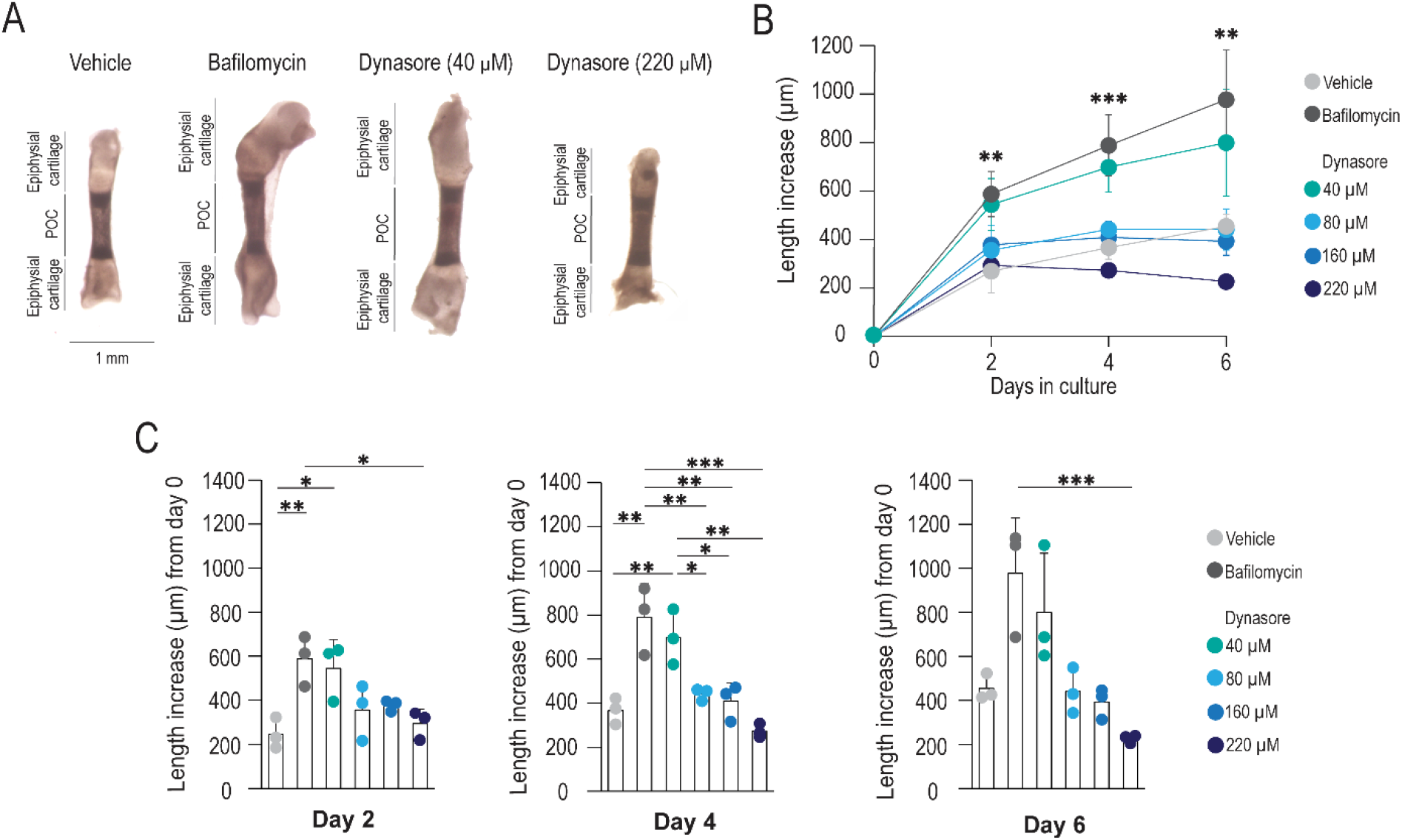
Dynasore leads to differential regulation on longitudinal bone growth. **A** *Ex vivo* cultures of metatarsals isolated from three-day-old mice were maintained in the presence of vehicle, bafilomycin (positive control) and dynasore (treatment). Representative images at day 6. **B, C** Longitudinal bone growth was assessed every two days. POC: primary ossification center. n = 3 animals (9 bones) for vehicle, bafilomycin and dynasore. Asterisks in panel B indicate statistically significant differences assessed by one-way ANOVA. Data is presented as mean ± SD. *p < 0.05. **p < 0.01. ***p < 0.001.

**Fig. 3.**
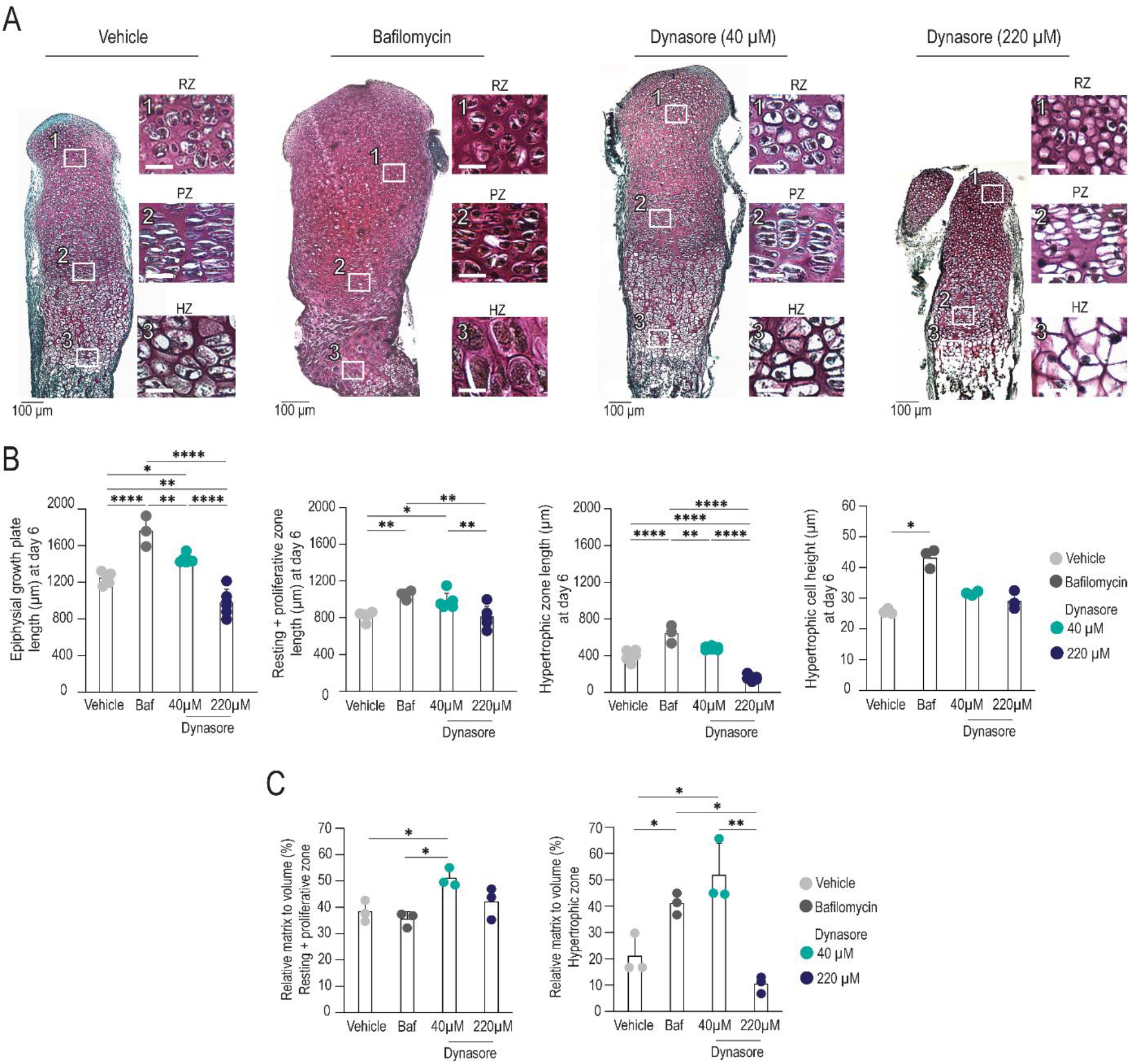
A low concentration of dynasore (40 µM) stimulates bone growth. **A** Histological characteristics of the epiphysial growth plate zones: resting, proliferative, and hypertrophic zones. **B** Bone growth measurements of the growth plate zones. **C** Relative matrix to volume assessment. Scale bar in insets = 25 µm. Baf: bafilomycin; HZ: hypertrophic zone; PZ: proliferative zones; RZ: resting zones. Data in A and B is presented as mean ± SD, n = 5 bones, except in bafilomycin and the hypertrophic cell height which represent 3 bones. *p < 0.05, **p < 0.01, and ****p < 0.0001.

Normality of the data was assessed using the Shapiro-Wilk test. When the assumption of normality was met, a statistical analysis was conducted using one-way ANOVA followed by Tukey’s multiple comparison test to evaluate differences between experimental groups, with significance levels denoted as *p < 0.05, **p < 0.01, ***p < 0.001, and ****p < 0.0001. When the assumption of normality was not met (Fig. 2C, day 6; Fig. 3B: hypertrophic cell height; Fig. 6B: resting, proliferative, and pre-hypertrophic zones), the Kruskal-Wallis test was used instead, followed by Dunn’s test for multiple comparisons, with the same significance thresholds. Graphs present data as mean ± SD, with individual data points included where appropriate. Details of data normalization are provided in the figure legends. All statistical analyses were performed using RStudio software (version 4.1.2, 2021-11-01) and graphs produced in Excel (2019 MSO. 16.0.10416.20047). Schematics were made with Biorender.com (agreement numbers: BY285AJ4ZS and ZV285AIZB6) and Illustrator (version 29.4. Adobe Creative Cloud).

## 3. Results

### 3.1. Dynasore has hormetic effects on bone elongation *ex vivo*

We used a well-established model to culture murine metatarsals *ex vivo* [12] and treated the bones with increasing concentrations of the inhibitor: 40 µM, 80 µM, 160 µM, and 220 µM (Fig. 1B, C). This dosage was based on previous studies reporting the use of dynasore, with concentrations of ≤ 80 µM typically achieving a strong inhibition of endocytosis [4, 7, 17]. Here, whilst the highest dose impaired normal growth, we were surprised to find that a dose of 40 µM stimulated longitudinal growth (Fig. 2B, C). Morphological changes caused by dynasore were evident from day 2 of culture, resulting in longer bones compared to both vehicle and higher concentrations (220 µM) by day 4 and 6 (Supplemental Fig. S1A, Fig. 2A-C). Of note, the lowest dose of dynasore (40 µM) supported robust bone growth to a similar extent as bafilomycin (Fig. 2A-C), a well-established promoter of metatarsal elongation [13]. On the contrary, 220 µM dynasore caused a marked reduction in bone length by the end of the culture period (Fig. 2A-C). These changes were also evident in the assessment of bone growth rate, which demonstrated that all tested dynasore concentrations initially promoted bone growth until day 2. However, in the subsequent days, the growth rate declined sharply across all conditions, except in bones treated with 40 µM dynasore, where longitudinal bone growth was sustained in a comparable manner to the bafilomycin treatment (Supplemental Fig. S1B).

### 3.2. Dynasore alters bone growth through differential dose-dependent effects on chondrocytes

To explore the changes induced by dynasore, we selected bones from the doses giving the strongest hormetic effects, 40 µM (which stimulated growth) and 220 µM (which repressed growth) for further investigation. Histological analysis of the bones treated with 40 µM dynasore revealed typical chondrocyte phenotype, with rounded cells in the resting zone (RZ) and columnar, flattened chondrocytes in the proliferative zone (PZ) (Fig. 3A), and enlarged cells in the hypertrophic zone (HZ) (Fig. 3A). By contrast, a high dose of dynasore (220 µM) led to empty lacunae in the resting and proliferative zones (Fig. 3A), suggesting potential cytotoxicity, which aligns with prior studies indicating that excessive disruption of dynamin activity can lead to cellular stress and apoptosis [18].

In order to assess changes in chondrocyte behaviour that could explain the increase in bone growth caused by the 40 µM dynasore, we conducted histomorphometry. Firstly, we verified that bone growth had increased due to the effects of dynasore on chondrocytes, since the length of the epiphyseal cartilage was significantly increased in the presence of 40 µM dynasore in comparison with vehicle (Fig. 3A). Interestingly, greater differences were observed in the resting/proliferating zones (Fig. 3B) than the hypertrophic zone height or in the height of the terminal hypertrophic cells (Fig. 3B). In contrast, bafilomycin stimulated elongation throughout the epiphyseal cartilage (Fig. 3B), whereas the reduced bone growth observed with the higher dose of dynasore (220 µM) was reflected in the reduced sizes of all parameters. To monitor changes in ECM turnover that could be mediated by dynamin-mediated endocytosis, we quantified the relative amount of ECM in the epiphysis. Interestingly, the resting, proliferating and hypertrophic zones contained relatively more ECM in bones treated with dynasore at 40 µM (Fig. 3C).

### 3.3. Low dose of dynasore decreased chondrocyte proliferation while maintaining collagen type X synthesis

To understand the cellular changes induced by dynasore, we labelled cells dividing in the last 4 hours of culture, using the thymidine analogue, EdU (Fig. 4A). Whereas vehicle-treated chondrocytes contained a relatively large proportion of dividing cells, all other treatments were associated with reduced cell division; this was most notable at the highest concentration (220 µM) of dynasore, which completely abolished chondrocyte proliferation as indicated by a total absence of EdU-positive cells (Fig. 4B, C). Even though the resting/proliferating zones were longer in dynasore 40 µM treatment, there was a reduced cell division rate in comparison to vehicle-treated bones. This was similar to bafilomycin treatment, which reduced cell proliferation despite increasing bone length (Fig. 4B, C) [13].

**Fig. 4.**
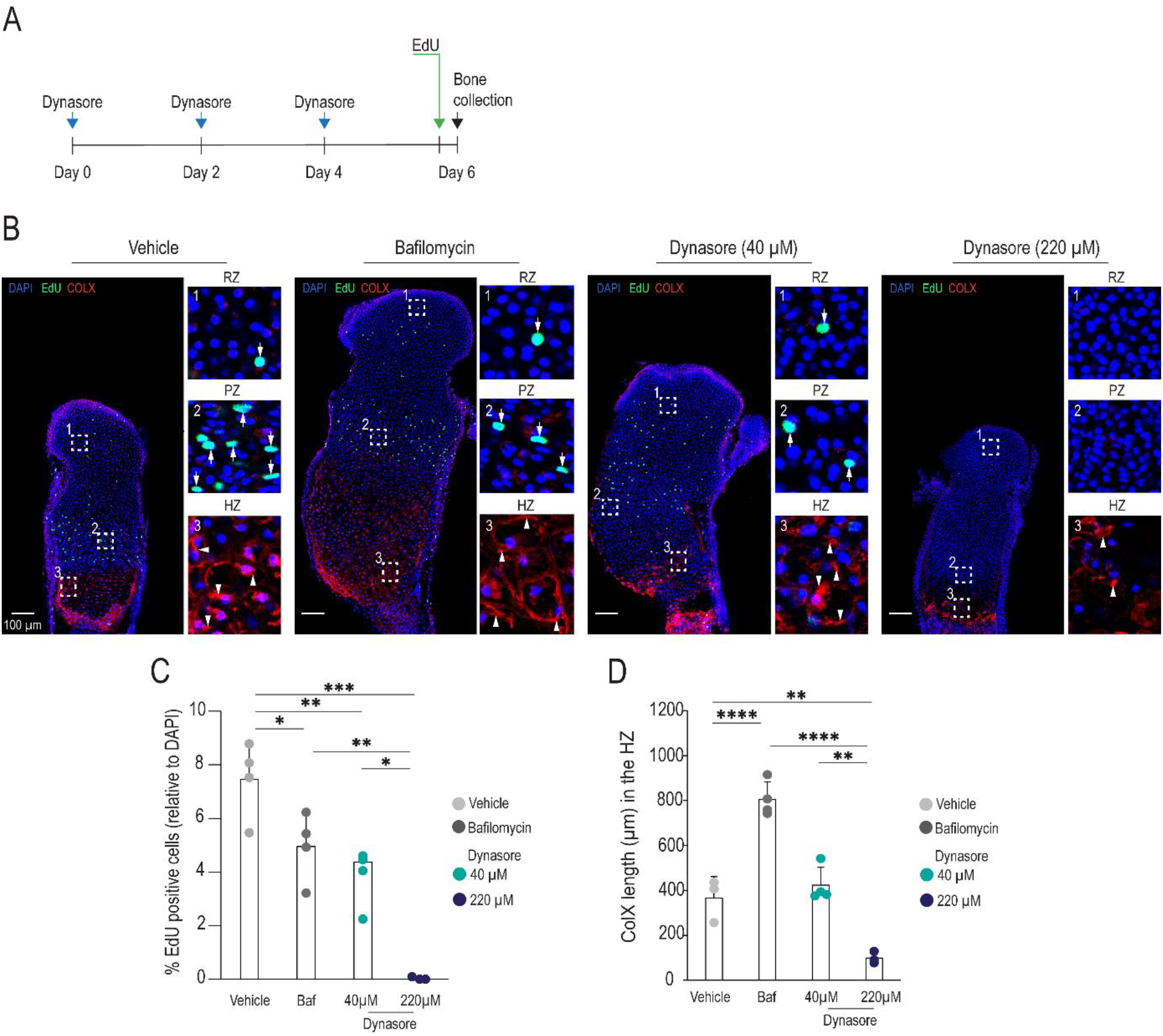
A low concentration of dynasore (40 µM) decreases chondrocyte proliferation while preserving collagen type X production. **A** Schematic representation of the bone cultures. EdU was added four hours before collecting the metatarsals at day 6. **B** EdU incorporation (white arrows in insets, visualised by Click assay) and collagen type X production (white arrowheads in insets, visualised by immunofluorescence) in chondrocytes. **C** Quantification of EdU positive chondrocytes and **D** the length of the collagen type X-positive region. Baf: bafilomycin; HZ: hypertrophic zone; PZ: proliferative zones; RZ: resting zones. n = 4 bones obtained from one litter, data is presented as mean ± SD. *p < 0.05, **p < 0.01, ***p < 0.01, and ****p < 0.0001.

To visualise the differences between proliferating and hypertrophic chondrocytes, we multiplexed the EdU staining protocol with collagen type X immunostaining (Fig. 4B, D). Type X collagen-stained ECM likely represents a combination of cartilage contained in the metatarsals at the point of tissue dissection as well as changes occurring during the culture period. Whilst bafilomycin stimulated the anticipated increase in chondrocyte hypertrophy in comparison to the vehicle-treatment [13], no significant changes were induced by dynasore at 40 µM. In contrast, exposure to 220 µM dynasore resulted in a marked reduction in collagen type X-stained region.

### 3.4. Dynasore, at low concentrations, blocks autophagy and activates mTORC1 pathway in growing metatarsal bones

One reported dynamin-independent target of dynasore is vacuolar-ATPase (v-ATPase), the specific target of bafilomycin. As there were similarities in metatarsal growth triggered by bafilomycin and dynasore (40 µM), we tested whether v-ATPase was inhibited in metatarsals following dynasore treatment. Since autophagy relies on lysosomal acidification by functional v-ATPase, the accumulation of the autophagy receptor, SQSTM1, can be used as a readout of impaired v-ATPase function [13]. Interestingly, we found that a similar percentage of chondrocytes were positive for SQSTM1 in the presence of 40 µM dynasore as with bafilomycin, which were both significantly elevated in comparison with the vehicle (Fig. 5A, B).

**Fig. 5.**
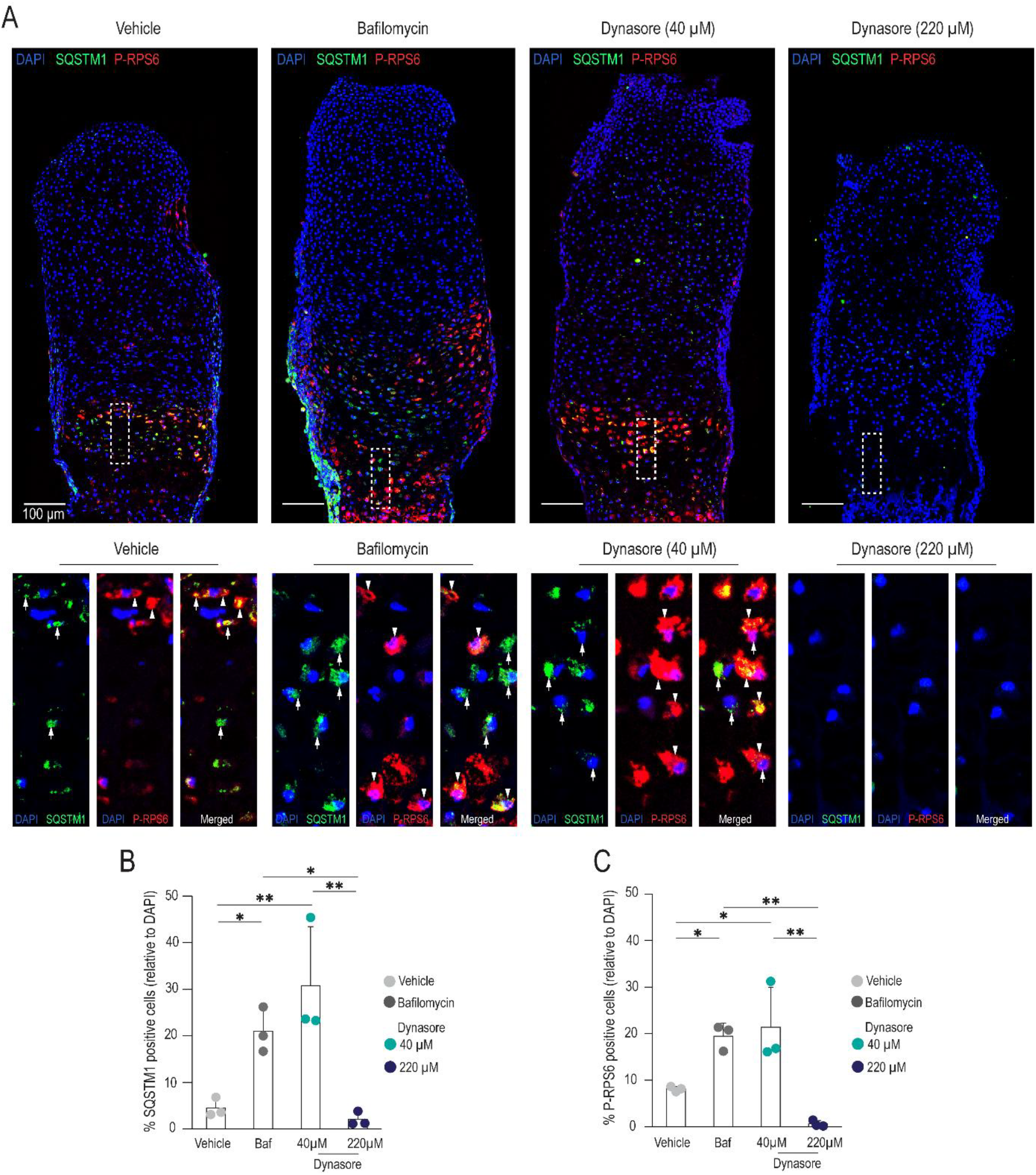
A low concentration of dynasore (40 µM) activates phosphorylated-RPS6 and accumulation of SQSTM1 in growing metatarsal bones. **A** Bones treated with 40 µM of dynasore showed higher accumulations of SQSTM1 (white arrowheads in insets) and increased levels of phosphorylated-RPS6 (white arrows in insets) in the growth plate, as visualised by immunofluorescence. **B, C** Quantifications of SQSTM1 and phosphorylated-RPS6. Baf: bafilomycin. Each data point was obtained from an independent experiment containing 3 bones per group; data is presented as mean ± SD. *p < 0.05, **p < 0.01.

Bafilomycin treatment also triggers activation of RPS6, a key readout of mTORC1 activity, in epiphyseal cartilage (Fig. 5A,C) [13]. We observed an increased phosphorylation of RPS6 in the pre-hypertrophic and hypertrophic zones at a low dose of dynasore (40 µM) (Fig. 5A-C). By contrast, both SQSTM1 and phosphorylated RPS6 were drastically decreased when higher doses of dynasore (220 µM) were used (Fig. 5A-C). These results indicate that 40 µM dynasore could inhibit the lysosomal v-ATPase function in chondrocytes.

### 3.5. Endocytosis of ECM is blocked by dynasore

While our results indicated that dynasore and bafilomycin both stimulated longitudinal growth, only some of the underlying mechanisms were similar (namely, decreased chondrocyte proliferation; phosphorylated RPS6 and SQSTM1 levels) whereas others were different: bafilomycin stimulated hypertrophy (based on histomorphometry and type X collagen immunostaining), whereas 40 µM dynasore led to no change in cell size but an increased relative amount of ECM. To test whether alterations in endocytosis of material from the ECM could explain these differences, we first assessed changes in filamentous actin (f-actin) distribution, which is typically enriched at the plasma membrane in growth plate chondrocytes [19], and included pre-HZ to assess the process of hypertrophy. Interestingly, we noted a wider layer of f-actin at the cell perimeter of dynasore-treated bones; this pattern was present in RZ (Fig. S2), PZ (Fig. S2) and pre-HZ chondrocytes (Fig. 6A), while no changes were observed in cells that were already hypertrophic (Fig. S2, Fig. 6B).

**Fig. 6.**
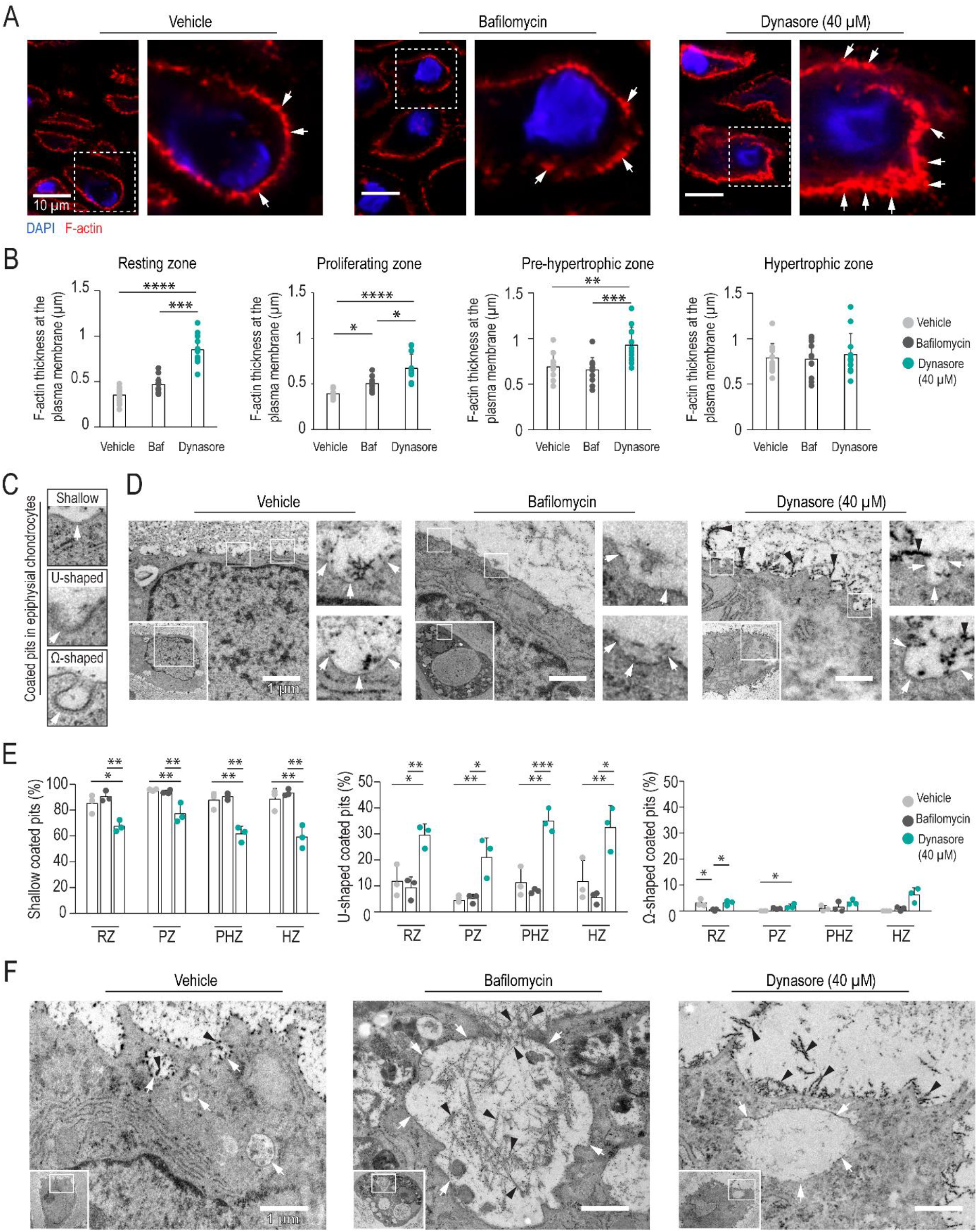
Endocytosis of cartilage matrix is impaired by dynasore. **A** - **B** Super-resolution microscopy images showing f-actin staining at the plasma membrane of pre-hypertrophic chondrocytes (white arrows) and quantification of the f-actin thickness in all growth plate zones. **C** Transmission electron microscopy (TEM) images of examples of shallow, U-shaped and Ω-shaped endocytic pits (white arrows) observed in the metatarsals. **D** Coated pits in pre-hypertrophic chondrocytes. Please, note the extracellular over-accumulation of matrix and collagen fibers (black arrow heads) in the dynasore-treated group. **E** Quantification of shallow, U-shaped and Ω-shaped pits in the epiphysial growth plate. **F** TEM images reveal cartilage endocytosis in hypertrophic chondrocytes. Chondrocytes uptake matrix and collagen fibers in endocytic pits/vesicles (white arrows), which were larger and contained more visible extracellular matrix in the bafilomycin treatment group (black arrowheads). By contrast, dynasore-treated bones show chondrocytes with over-accumulation of ECM in the extracellular space (black arrowheads). Each data point in B represent one cell whereas in E each data point is the average of three cells. All experiments are derived from 3 bones obtained from three different litters; data is presented as mean ± SD. *p < 0.05, **p < 0.01, and ****p < 0.0001.

To verify if this disrupted f-actin distribution may reflect altered membrane trafficking, we visualised the ultrastructure of cell membranes using transmission electron microscopy (TEM). While endocytic pits (Fig. 6C) were evident on chondrocytes in the RZ (Fig. S3), PZ (Fig. S3), pre-HZ (Fig. 6D) and HZ (Fig. S2C), in all conditions, dynasore altered pit morphology: whereas shallower pits were predominantly found in vehicle and bafilomycin-treated groups, dynasore caused pits to become deeper (Fig. 6E). This observation is consistent with the role of dynamin in vesicle scission, and disruption of this process by dynasore preventing internalization of the material [4]. Furthermore, we also observed that dynasore-treatment caused endocytic pits to be coated with a more electron dense material (Fig. 6D). We used the TEM images to screen endosomal compartments for evidence of internalised ECM. Structures resembling collagen fibers were evident, particularly in bafilomycin-treated group (Fig. 6F).

Together, these data suggest that while bafilomycin promotes bone elongation via chondrocyte hypertrophy (via blocking lysosomal v-ATPase, which activates mTORC1 signalling and the endosomal accumulation of endocytosed material due to impaired autophagy), dynasore stimulates longitudinal growth without promoting hypertrophy, which, despite also blocking lysosomal v-ATPase, appears to be caused by an accumulation of ECM in the extracellular space due to impaired cartilage remodelling.

To understand if these effects of dynasore and bafilomycin were mutually exclusive, we assessed the effect of co-treatment on metatarsal growth (Fig. S4). However, the simultaneous administration of dynasore and bafilomycin did not produce an additive or synergistic increase in bone length (Fig. S4 B-D). When applied alone, higher doses of both dynasore (Fig. 2B) and bafilomycin [20] cause growth impairments, and we suspect this effect may have occurred when these doses are combined, perhaps via further v-ATPase inhibition.

In summary, our findings indicate that at relatively low doses, prolonged use of dynasore can stimulate chondrocytes to cause tissue-wide effects that facilitated bone elongation, whereas at higher concentrations it has deleterious effects, typical of hormesis.

## 4. Discussion

As a small molecule, dynasore was identified as a potent inhibitor of the GTPase activity of dynamin-1 [4, 17] and subsequently was shown to potently inhibit dynamin function. However, further investigations revealed that it elicits effects in cells lacking dynamin genes [18] indicating dynamin-independent activities including altered cell membrane remodelling, destabilization of f-actin and v-ATPase inhibition [21]. Altogether, the use of dynasore to block dynamin becomes complicated by these multiple potential routes of action, which can be simultaneously affected and vary by cell type/state.

Using bafilomycin as the positive control for bone growth led us to directly compare the growth mechanisms with those of dynasore, which we thought could be similar as both are able to inhibit the v-ATPase. While we found some similarities with metatarsal growth induced by bafilomycin [13], specifically an increase in size of RZ+PZ length, reduced proliferation and elevated phosphorylated RPS6 and SQSTM1 accumulation, phenotypes were not identical since there was no marked increase in hypertrophic zone parameters measured either by histology or collagen type X staining in the presence of dynasore. By exploring cellular ultrastructures, we were able to identify collagen fibers within endosomal compartments in the metatarsals. Since collagen fibers are assembled in the extracellular space [22], it is highly likely that this material was endocytosed rather than being retained in the cell via, for example, impaired exocytosis; the fact that collagen fibers were most evident in the bafilomycin-treated group may be due to the actions of this bafilomycin to block their degradation via autophagy, causing them to accumulate. This endocytosed material may contribute to the dry mass accumulation during hypertrophy of the chondrocytes [23], suggesting that dynamin-mediated endocytosis is needed for ECM remodelling and chondrocyte hypertrophy.

Although we focused on the growth-promoting effects of dynasore at a dose of 40 µM, our results showed a hormetic effect: whereas low doses had a growth promoting effect, higher doses inhibited growth [6]. Hormetic responses, in which opposing effects occur at high and low doses of the same stimuli, are well described in the literature [6]. Given that dynasore has been shown to induce cell death [24], and nuclear abnormalities and empty lacunae were visible following six days with 220 µM dynasore treatment, dosing must be carefully considered. It was interesting to note that at doses safe to use in other tissues [7], 80 µM, for example, bone growth rates were very low after an initial stimulation of growth by day 2 of treatment.

Interestingly, dynasore may be safe to use to target chondrocytes *in vivo*. Indeed, dynasore administration to mice by intra-articular injection during osteoarthritis development, aiming to target articular chondrocytes, preserved cartilage integrity: fortnightly dynasore injections could reduce cartilage destruction (OARSI score) caused by destabilization of the medial meniscus (DMM) surgery in both wild-type mice and those with inducible chondrocyte-specific myosin light chain 3 mutations. The dynasore-induced amelioration was associated with a reduction in senescent markers p16^INK4A^ and γH2AX brought about by OA development, which was attributed to reduced cartilage degradation via clathrin-mediated endocytosis [9]. The clear positive effects of treatment are interesting given the intermittent application of dynasore in this experiment, consistent with its application in other studies [25]. In the case that dynasore may be applied *in vivo* to test its ability to stimulate bone growth, establishing the optimal dose will be important to avoid the negative effects of over-exposure. Furthermore, long-term exposure must be checked since inhibiting cartilage endocytosis and turnover would inevitably impair continued growth. Finding precisely how dynasore stimulates growth would be preferable, however, to allow more specific small molecules to be used and mitigate unwanted effects. Hence, dynasore is a valuable tool for investigating dynamin-mediated processes like endocytosis, but like all inhibitors, its use requires caution. Understanding its mechanisms of action, optimal concentration range, and potential off-target effects will be crucial for drawing accurate conclusions about its hormetic role in bone growth.

## Conclusions

In summary, our findings suggest that the dynamin inhibitor, dynasore, promotes longitudinal bone growth at low concentrations, whilst at higher concentrations it exerts deleterious effects.

## Acknowledgements

This work was supported financially by the Novo Nordisk Foundation (Phillip T. Newton: 0067241), Swedish Research Council (Phillip T. Newton: 2019-01919; Oscar P.B. Wiklander: 2022-02449), CIMED (Oscar P.B.Wiklander: FoUI-976434) and Karolinska Institutet (Phillip T. Newton; Oscar P.B. Wiklander). We also wish to thank the Biomedicum imaging core facility (BIC), HistoCore – Biomedicum, Preparative Histocore, comparative medicine biomedicum (KM-B) and the electron microscopy unit Emil at Karolinska Institutet for their support.

## Conflict of interest statement

Oscar P.B. Wiklander has a financial interest in Evox Therapeutics.

## Data availability

The underlying data for this study are available upon request.

