## Supplementary Figures 1-4 for "Dynasore, the dynamin inhibitor, modulates longitudinal bone growth in a hormetic manner"

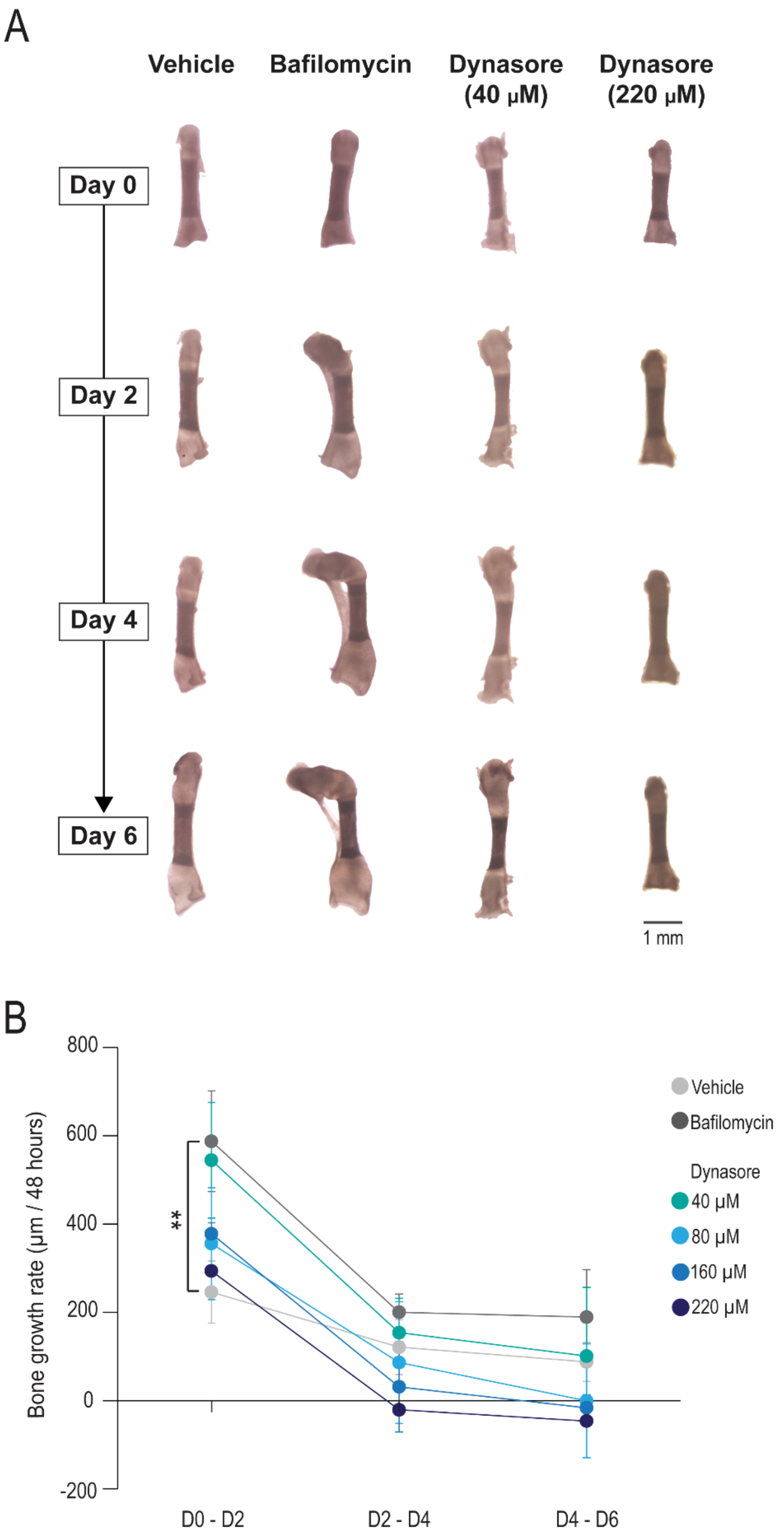


**Figure supplementary 1.** Longitudinal bone growth of metatarsal *ex vivo* cultures from day 0 to day 6. **A** Microphotographs of metatarsal bones were collected during culture. **B** Bone growth rate. Data show three independent experiments per group and is presented as mean ± SD. $^{**}p < 0.01$.

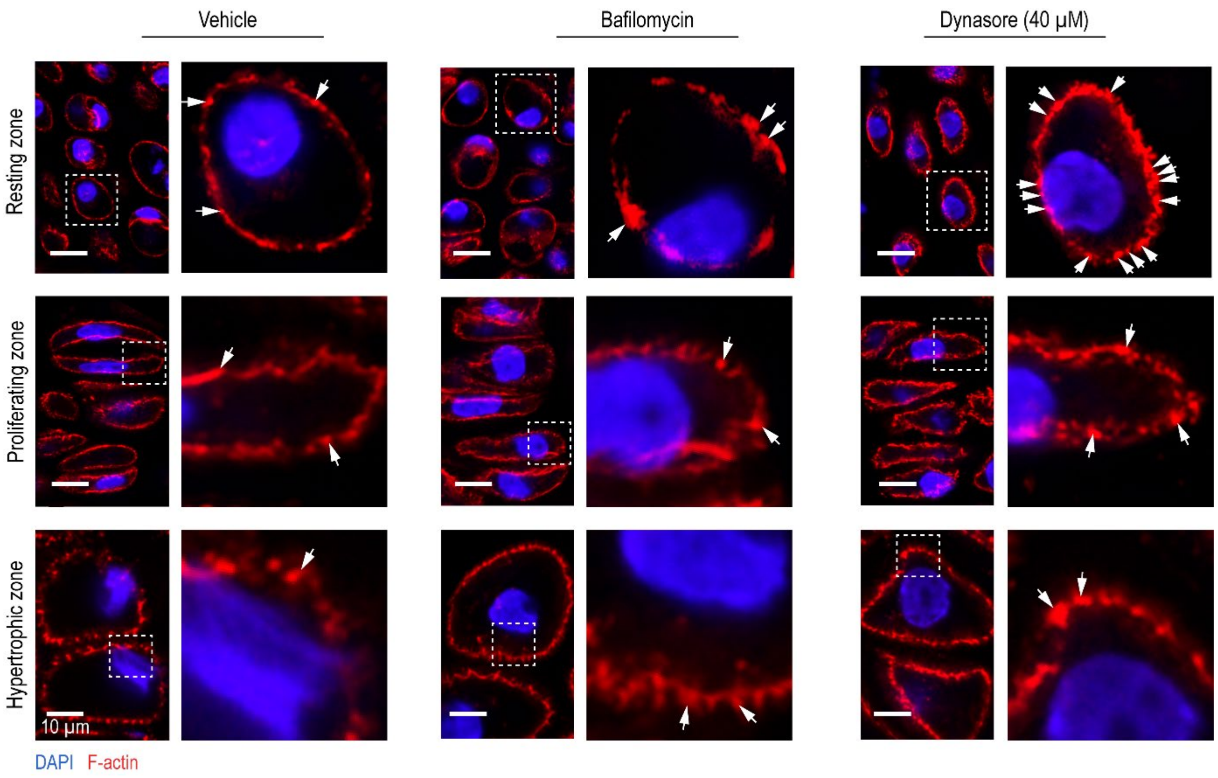


**Figure supplementary 2.** Super-resolution microscopy images showing f-actin staining (white arrows) at the plasma membrane of growth plate chondrocytes. N = 3 bones from three different litters.

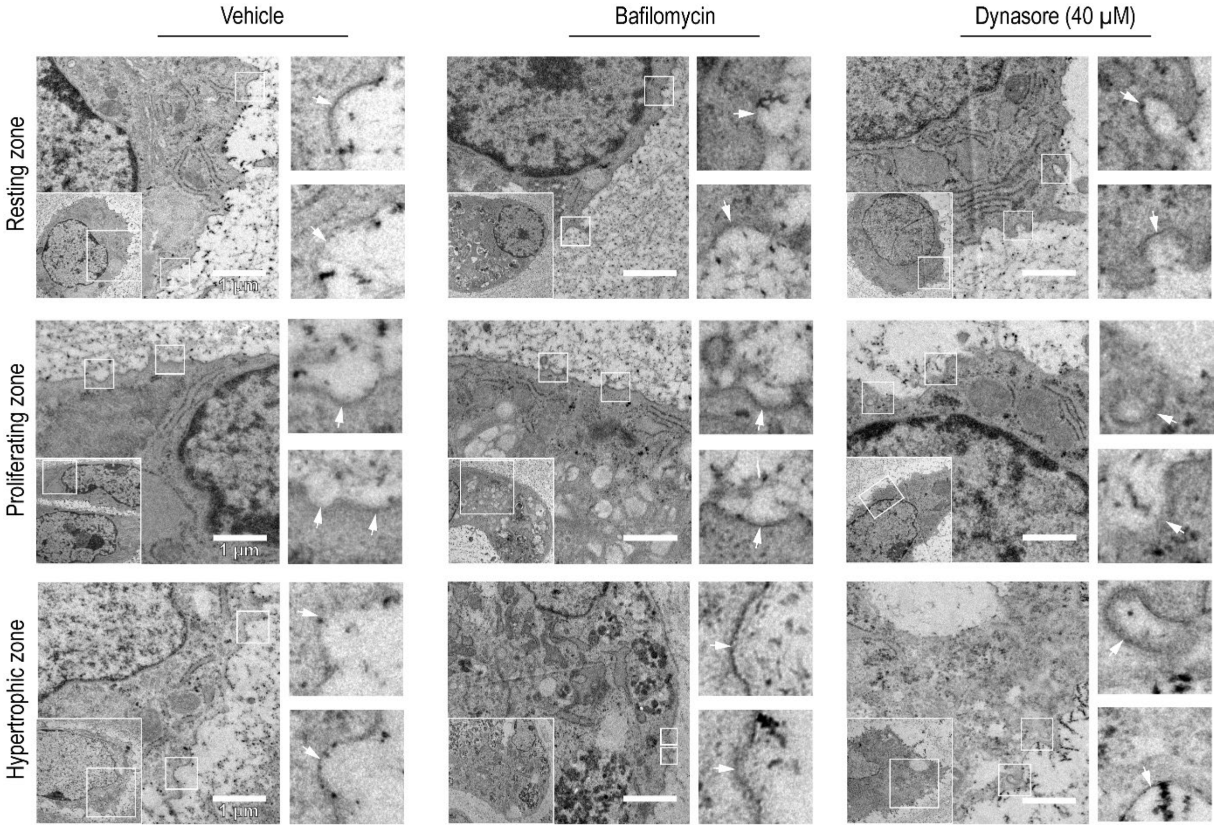


**Figure supplementary 3.** Transmission electron microscopy (TEM) images of growth plate chondrocytes to visualise endocytic pits (white arrows). All experiments are derived from 3 bones obtained from three different litters.

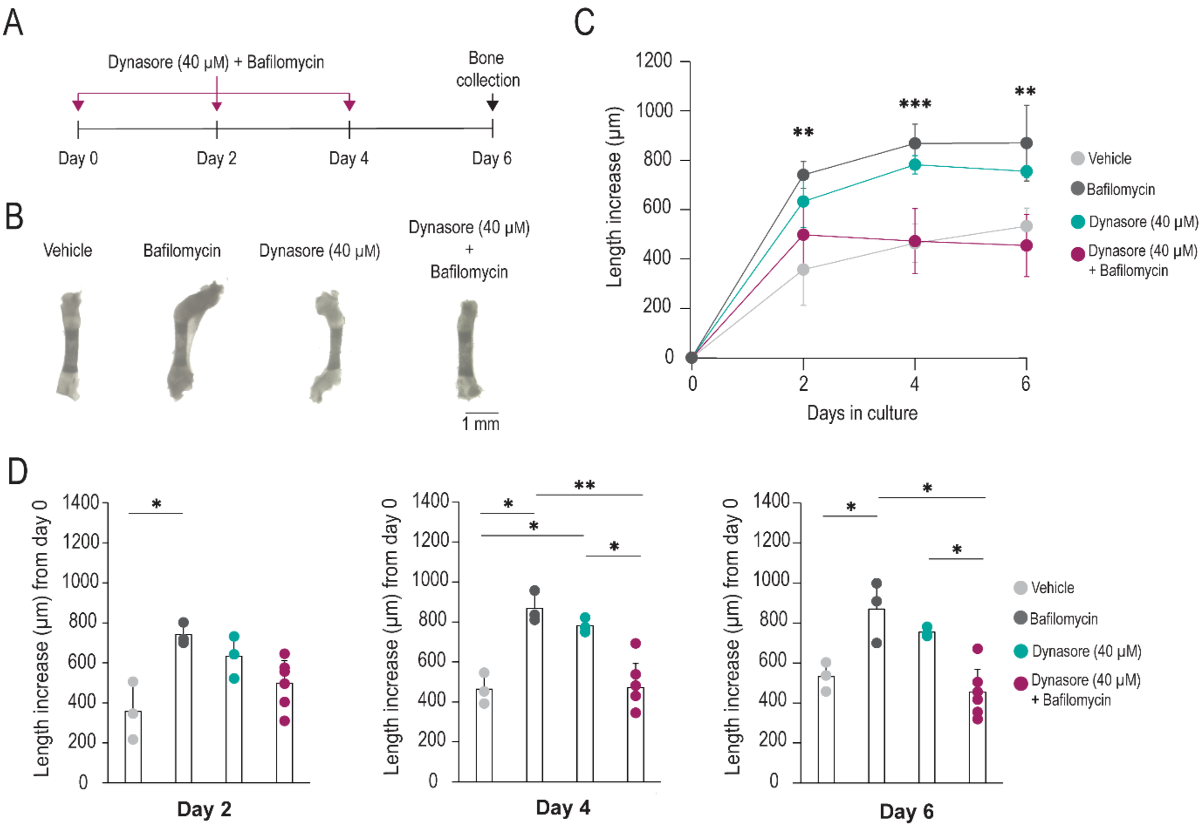


**Figure supplementary 4.** Bafilomycin and dynasore (40 μM) co-administration does not additively or synergistically enhance bone growth. **A** schematic of the experimental approach. **B** Representative images at day 6 of metatarsals *ex vivo* cultures. Bones were maintained in the presence of vehicle, bafilomycin, dynasore (40 μM), and dynasore (40 μM) + bafilomycin (treatment). **C, D** Longitudinal bone growth was assessed every two days. *n* = 3 bones for vehicle, bafilomycin and dynasore (40 μM), except for dynasore (40 μM) + bafilomycin which has an *n* = 6 bones. Data is presented as mean ± SD. *$p < 0.05$. **$p < 0.01$.
